# Inducible, tunable and multiplex human gene regulation using CRISPR-Cpf1-based transcription factors

**DOI:** 10.1101/150656

**Authors:** Yu Gyoung Tak, Benjamin P. Kleinstiver, James K. Nuñez, Jonathan Y. Hsu, Jingyi Gong, Jonathan S. Weissman, J. Keith Joung

## Abstract

Targeted and inducible regulation of mammalian gene expression is a broadly important research capability that may also enable development of novel therapeutics for treating human diseases. Here we demonstrate that a catalytically inactive RNA-guided CRISPR-Cpf1 nuclease fused to transcriptional activation domains can up-regulate endogenous human gene expression. We engineered drug-inducible Cpf1-based activators and show how this system can be used to tune the regulation of endogenous gene transcription in human cells. Leveraging the simpler multiplex capability of the Cpf1 platform, we show that we can induce both synergistic and combinatorial gene expression in human cells. Our work should enable the creation of other Cpf1-based gene regulatory fusion proteins and the development of multiplex gene perturbation library screens for understanding complex cellular phenotypes.

---

Sequence-specific RNA-guided CRISPR-Cas nucleases have revolutionized biological research due to their ease of programmability. The widely used Cas9 from *Streptococcus pyogenes* (**SpCas9**) can be targeted to specific DNA sequences by an associated complementary guide RNA (**gRNA**), provided that a protospacer adjacent motif (**PAM**) of the form NGG is also present. Catalytically inactive SpCas9 (“dead” SpCas9 or **dSpCas9**) have been fused to transcriptional activation or repression domains to alter the expression of individual genes^1, 2^ or to perform genome-wide library screens^3-5^ in mammalian cells. Both small molecule- and light-inducible dSpCas9-based fusions have been developed^6-10^, enabling the ability to regulate the activity of this platform. Recently described CRISPR-Cpf1 nucleases offer additional capabilities beyond those of SpCas9 including shorter length CRISPR RNAs (**crRNA**s) for guiding Cpf1 to targets, the ability to target T-rich PAMs^11-13^, and RNase processing of multiple crRNAs from a single transcript by sequence-specific ribonuclease activity resident within Cpf1 itself^14, 15^. However, to our knowledge, “dead” Cpf1-based gene regulators have thus far only been shown to repress gene expression in bacteria^16, 17^ and plant (*Arabidopsis*)^18^ and have not been shown to function in mammalian cells either as activators or repressors. Here we describe both constitutively active and chemically inducible Cpf1-based transcriptional activator platforms for upregulating human gene promoters. We also demonstrate that facile expression of multiple crRNAs in a single transcript can be leveraged to achieve synergistic or combinatorial activation of endogenous genes in human cells.

We first tested whether we could upregulate endogenous human gene promoters with direct fusions of catalytically inactive Cpf1 nuclease to the strong synthetic **VPR** activator, which consists of four copies of the Herpes Simplex Virus-derived VP16 activator, the human NF-KB p65 activation domain, and the Epstein-Barr Virus-derived R transactivator (Rta)^19^ (**Fig. 1a**). Initial GFP reporter-based assay experiments showed that a “dead” Cpf1 from *Lachnospiraceae bacterium ND2006* (**dLbCpf1**)-VPR fusion induced higher levels of gene activation than a “dead” Cpf1 from *Acidaminococcus sp. BV3L6* (**dAsCpf1**)-VPR fusion (**Supplementary Fig. 1**). This difference is consistent with previous reports from our group and others showing that LbCpf1 generally exhibits higher genome-editing activities than AsCpf1 in human cells^20, 21^. Therefore, for all subsequent experiments described herein, we used the dLbCpf1 protein.

**Figure 1.**
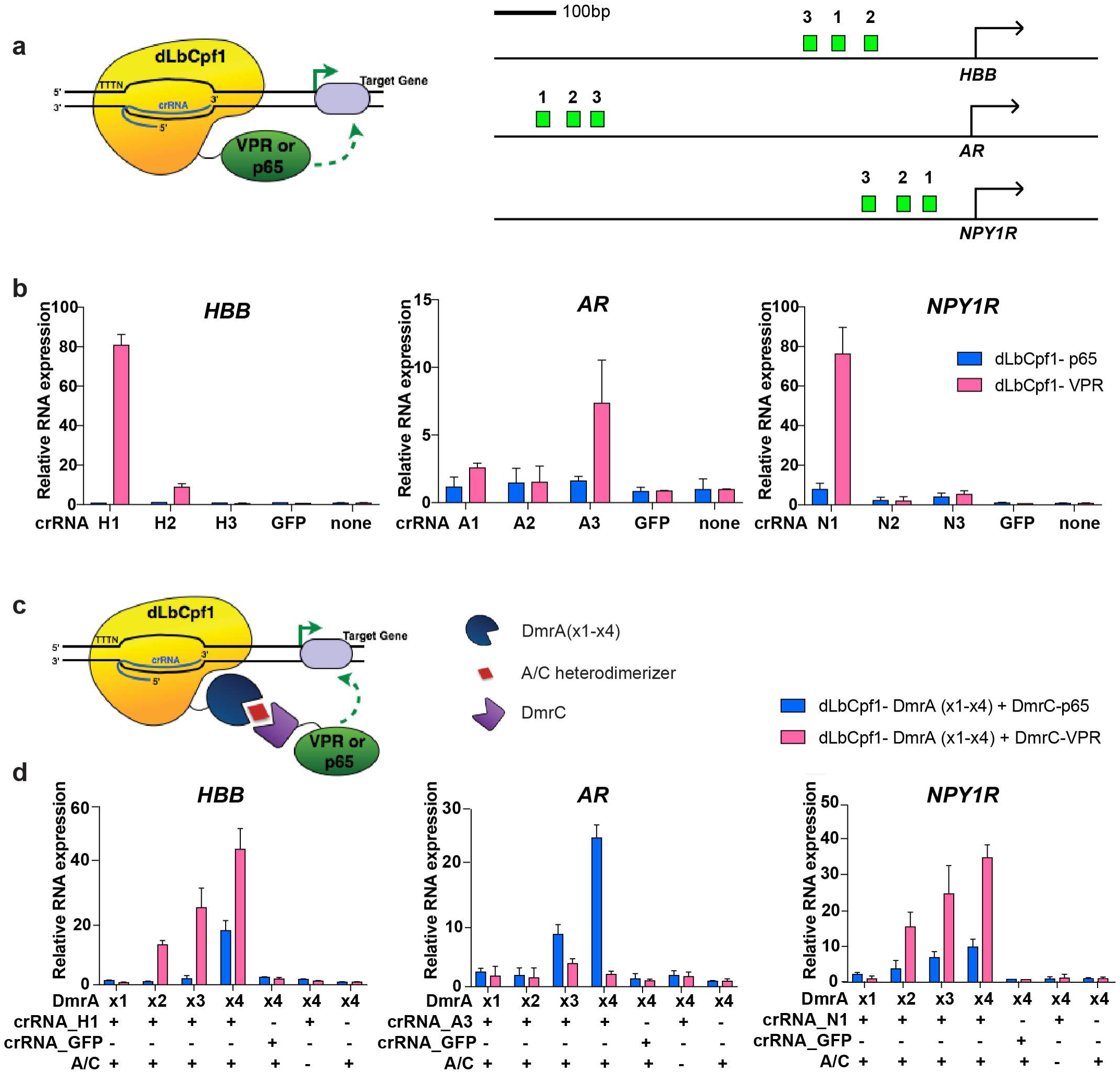
Targeted human endogenous gene regulation using individual crRNAs with dLbCpf1-based activators. a. Schematic showing direct dLbCpf1-VPR and dLbCpf-p65 activator fusion proteins. The binding sites for crRNAs (green boxes) targeting promoters of each gene are shown on the right panel.
b. Activities of dLbCpf1-p65 or dLbCpf1-VPR using single crRNAs at three endogenous human genes (*HBB*, *AR*, and *NPY1R*) in HEK293 cells. Relative activation of the indicated gene promoters was measured by RT-qPCR. Three separate individual crRNAs were targeted within promoter sequence 1 kb upstream of the transcription start site for each gene. Relative mRNA expression is calculated by comparison to the control sample in which no crRNA is expressed.
c. Schematic illustrating drug-dependent bi-partite dLbCpf1-based activator fusion proteins. dLbCpf1 is fused to one to four DmrA domains and VPR or p65 is fused to a DmrC domain. Because DmrA and DmrC interact only in the presence of a A/C-heterodimerizer drug (red diamond), the bi-partite activator is only reconstituted in the presence of the drug.
d. Activities of drug-dependent bi-partite dLbCpf1-based activators using single crRNAs at three endogenous human genes (*HBB*, *AR*, and *NPY1R*) in HEK293 cells. Relative activation of the indicated gene promoters was measured by RT-qPCR. The crRNA used for each promoters was the one that showed the highest activity in (b) above. Representative data shown in **(b)** and **(d)** are the means of three biological independent replicates and error bars indicate standard deviation (SD).

We next targeted the promoters of three different endogenous genes that are expressed at low levels in human HEK293 cells (*HBB*, *AR* and *NPY1R*) by designing three crRNAs for each promoter, located at various distances upstream of the transcription start point (**Fig. 1a**). Testing each of these crRNAs with the dLbCpf1-VPR fusion demonstrated robust transcriptional activation with at least one crRNA for each of the three target genes as assayed by measurement of transcription by real-time RT-PCR (**Fig. 1b**). Testing of these same crRNAs with direct fusions of dLbCpf1 to the NF-KB p65 activation domain alone resulted in either little or no transcriptional activation of the target gene promoter (**Fig. 1b**). Thus, as has been previously observed with dSpCas9-based activators, upregulation of gene expression by direct dLbCpf1 fusions is more efficient when multiple strong activation domains are recruited to a target promoter^3, 5, 19, 22^.

To extend the utility of dLbCpf1-based activators, we sought to construct drug-regulated versions of these effectors. To do this, we used the well-characterized DmrA and DmrC domains (fragments of the FK506-binding protein (FKBP) and FKBP-rapamycin-binding protein (FRB), respectively) that interact only in the presence of a rapamycin analog known as the A/C heterodimerizer^6, 23, 24^. We envisioned creating split dLbCpf1 activators consisting of two fusions: dLbCpf1 fused to a DmrA domain and a DmrC domain fused to VPR (**Fig. 1c**). In this configuration, a reconstituted activator should assemble only in the presence of the A/C drug. Testing of these bipartite fusions in the presence of the A/C heterodimerizer with single crRNAs (shown above to work efficiently with the direct dLbCpf1 activator fusions) failed to reveal activation of the *HBB*, *AR* or *NPY1R* genes (**Fig. 1d**). We reasoned that increasing the number of DmrA domains linked to dLbCpf1 (and therefore the number of VPR domains recruited to the promoter) might increase the efficiency of gene activation. Testing of dLbCpf1 fusions harboring two, three or four tandem copies of the DmrA domain in the presence of DmrC-VPR revealed activation at two of the three endogenous gene promoters we tested (*HBB* and *NPY1R*; **Fig. 1d**). Activation levels observed correlated with the number of DmrA domains, with maximum levels reaching approximately half of that observed with direct dLbCpf1-VPR fusions. As expected, activation was contingent on the presence of the A/C heterodimerizer drug (**Fig. 1d**), demonstrating drug-dependency of this system.

Surprisingly, we found that using DmrC-p65 fusions with dLbCpf1-DmrA fusions led to drug-dependent transcriptional upregulation from all three target gene promoters (**Fig. 1d**), an unexpected result given the lack of activation observed with direct dLbCpf1-p65 fusions and these same crRNAs (**Fig. 1b**). For *HBB* and *NPY1R*, the maximal levels of activation observed with DmrC-p65 were somewhat less (~50% and ~30%, respectively) than those obtained with DmrC-VPR but still robust in absolute terms (~20-fold and ~10-fold activation). For *AR*, DmrCp65 activated transcription by ~25-fold, showing superior activity relative to the lack of an effect by DmrC-VPR using the same crRNA.

One major advantage of Cpf1 compared to Cas9 is the ability to express multiple crRNAs within a single transcript that is subsequently processed into individual crRNAs by the RNase processing activity of Cpf1 (**Fig. 2a**), thereby better enabling multiplex applications^14, 15, 17^. Expressing multiple gRNAs for SpCas9 from separate promoters can be challenging due to substantial recombination that can occur between promoter sequences and the crRNA:tracrRNA constant regions^25, 26^ or the need for additional accessory RNA sequences or trans-acting factors if multiple gRNAs are expressed from a single transcript^26-32^. With Cpf1, multiple crRNAs can be encoded in one transcript, a capability that has been used previously to achieve multiplex gene editing in human cells^15^.

**Figure 2.**
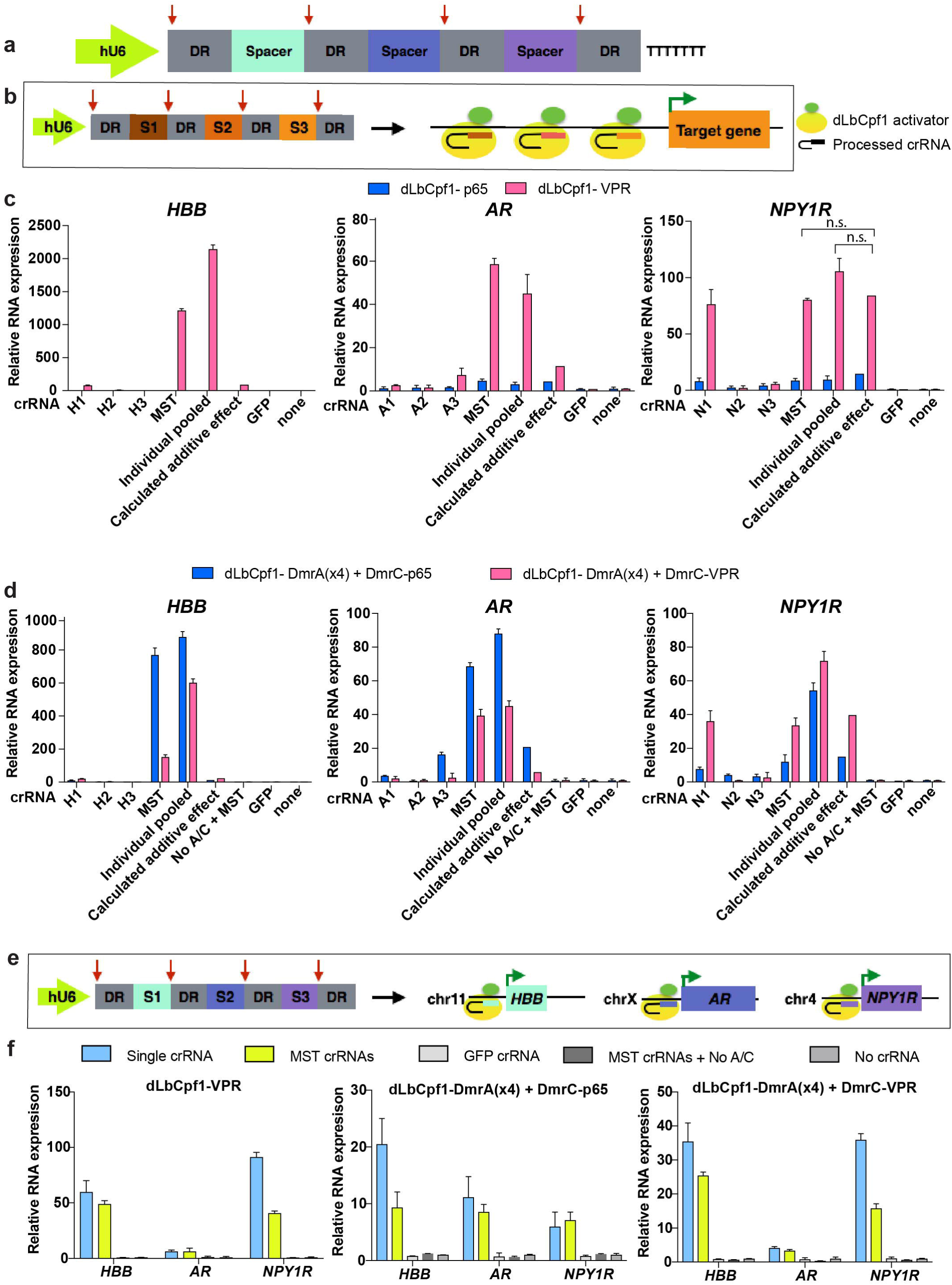
Multiplex and synergistic regulation of endogenous human genes by dLbCpf1-based activators. a. Schematic of an expression cassette designed to express multiple gRNAs encoded on a single transcript. The arrows (red) indicate cleavage sites being processed by the RNase activity of dLbCpf1.
b. Schematic illustrating multiplex expression of three crRNAs each targeted to the same endogenous gene promoter in the same cell.
c. Activities of direct dLbCpf1-p65 or dLbCpf1-VPR fusions with sets of three crRNAs expressed from a multiplex transcript or from individual transcripts on the HBB, AR, or NPY1R endogenous gene promoters. Transcripts were measured in HEK293 cells using RT-qPCR with relative mRNA expression calculated by comparison to the control sample in which no crRNA is expressed.
d. Activities of dLbCpf1-DmrA(x4) and DmrC-VPR fusions (pink bars) or with dLbCpf1-DmrA(x4) and DmrC-p65 fusions (blue bars) with sets of three crRNAs expressed from a multiplex transcript or from individual transcripts on the HBB, AR, or NPY1R endogenous gene promoters.
e. Schematic illustrating multiplex expression of three crRNAs each targeted to a different endogenous gene promoter in a single cell.
f. Simultaneous activation of three endogenous human genes using crRNAs expressed from a multiplex transcript or from individual transcripts with dLbCpf1-VPR direct fusions (left panel), dLbCpf1-DmrA(x4) and DmrC-VPR fusions (middle panel), and dLbCpf1-DmrA(x4) and DmrCp65 fusions (right panel). Transcripts were measured in HEK293 cells using RT-qPCR with relative mRNA expression calculated by comparison to the control sample in which no crRNA is expressed. Representative data shown in **(c), (d)** and **(f)** the means of three biological independent replicates and error bars indicate standard deviation (SD). hU6, human U6 Polymerase III promoter; DR; direct repeat sequence; n.s., not significant by Student t-test (p > 0.05)

To test whether multiplex single transcript (**MST**) crRNAs might be used with dLbCpf1-based activators, we used the same sets of crRNAs we designed above for the *HBB*, *AR* or *NPY1R* genes and encoded sets of three crRNAs directed against a single promoter in one transcript (**Fig. 2b**). We reasoned that if a set of MST crRNAs were active in the same cell, then we might observe synergistic increases in transcription from the target gene promoter (**Fig. 2b**), with synergy defined as greater than additive effects induced by two or more activators acting on the same promoter^2^. With direct fusions of VPR to dLbCpf1, we observed synergistic activation for two of the three endogenous human gene promoters (*HBB* and *AR*; **Fig. 2c**). For the third gene promoter (*NPY1R*), we did not observe synergistic activation (**Fig. 2c**). We also failed to observe synergistic activation of any of the three genes with direct fusion of p65 to dLbCpf1, with either MST crRNAs or separate crRNAs (**Fig. 2c**), but this was not unexpected since none of these crRNAs individually activate transcription with dLbCpf1-p65 (**Fig. 1b**). These latter crRNAs might not be optimal for Cpf1 activation and the characteristics that define a robust crRNA remain to be elucidated in future experiments.

We also observed drug-dependent synergy with dLbCpf1-DmrA(x4) and DmrC-VPR at the *HBB* and *AR* promoters with MST crRNAs. As with the direct activator fusions, synergistic activation was not observed at the NPY1R promoter with MST crRNAs even though it was seen with individually expressed crRNAs introduced together (**Fig. 2d**). A similar pattern of results for these three gene promoters was observed with dLbCpf1-DmrA(x4) and DmrC-p65 (**Fig. 2d**). The synergistic activation levels observed with DmrC-p65 on the *HBB* and *AR* genes were higher than comparable experiments performed with DmrC-VPR (**Fig. 2d**).

Having established that multiple active crRNAs can be expressed from a single transcript, we next sought to determine whether MST crRNAs might be used to simultaneously up-regulate different endogenous human gene promoters in the same cells. To test this, we encoded three crRNAs, each already shown to robustly activate a different gene promoter with dLbCpf1-activators on a single transcript (**Fig. 2e**). We found that these MST crRNAs could be used together with (i) dLbCpf1-VPR direct fusions, (ii) dLbCpf1-DmrA(x4) and DmrC-VPR fusions, or (iii) dLbCpf1-DmrA(x4) and DmrC-p65 fusions to simultaneously mediate transcriptional activation of the endogenous *HBB*, *AR*, and *NPY1R* gene promoters in human HEK293 cells (**Fig. 2f**). The magnitudes of activation observed with the MST crRNAs were somewhat lower (~18 to 55%) compared with the same crRNAs expressed from separate expression vectors, suggesting that further optimization might be achieved by varying MST crRNA transcript architecture and/or protein and RNA stoichiometries. Nonetheless, these results demonstrate that as many as three crRNAs encoded in a single transcript can be used to simultaneously up-regulate transcription of multiple gene promoters using dLbCpf1-based activators.

To our knowledge, the results described here provide the first demonstration that Cpf1-derived proteins can be used regulate endogenous gene expression in human cells. The orthogonal PAM recognition specificities of wild-type LbCpf1 (TTTV) and two engineered Cpf1 variants^13^ (TYCV and TATV) will enable an increased range of targetable sequences relative to what can be achieved with SpCas9 (NGG), a capability that should be particularly useful for genome-wide screens with RNA-guided gene regulatory proteins. Our success in using dCpf1 proteins to target two different activation domains (VPR and p65) should also motivate the engineering of dCpf1-based fusions bearing other heterologous regulatory domains (e.g., p300, DNMT, TET1) as has been previously done with dSpCas9^33, 34^ Given that the genome-wide specificities of LbCpf1 nuclease are reported to be at least comparable (if not actually superior) to those of SpCas9 nuclease^20, 21^, it seems likely that dLbCpf1 activators will be as specific as dSpCas9 activators, which have been shown to generally have minimal or no off-target gene activation events^22, 33^; however, additional future experiments will be needed to confirm this.

Our work defines generalizable strategies for regulating and tuning the activities of RNA-guided gene activator proteins. The bipartite activators we engineered enabled a useful on-off capability that responds to the cell-permeable A/C dimerizer drug, similar to other systems that have been described^10, 35^. In addition, increasing the number of DmrA dimerizer domains fused to dLbCpf1 enabled tuning of activation levels, presumably by increasing the number of DmrCactivator domain fusions recruited. These strategies could be easily extended to other functional domains beyond p65 and VPR and to more RNA-guided proteins such as other Cpf1 and Cas9 orthologues. In this regard, we note that we have successfully used these same strategies to regulate and tune dSpCas9-based activators (**Supplementary Fig. 2**) as have others who published similar drug-regulated dSpCas9-based activators while this work was in progress^8^.

We have also demonstrated that a key advantage of the Cpf1 platform, the ability to more easily encode multiple crRNAs on a single transcript, can be leveraged to enable synergistic or multiplex gene activation in human cells. Synergy with multiple crRNAs targeted to the same promoter could be used to increase the magnitude and success rate of target gene activation in library screens. Multiplex gene activation with crRNAs directed to different promoters could potentially be exploited to perform more complex library screens in which the expression of two or more genes are simultaneously altered, thereby enabling analysis of more complex cellular phenotypes with a multi-gene perturbation approach. Because up to three crRNAs can be readily fit on an oligonucleotide made by chip-based synthesis, precisely defined crRNA combinations of interest can be constructed for genes of interest, thereby more efficiently managing the size of libraries compared with library construction strategies that rely on random combinations of crRNA pairs. Overall, our findings should enable and motivate the use of dCpf1-based gene regulatory proteins for performing both focused and genome-wide gene perturbation screens in mammalian cells.

## ONLINE METHODS

### Plasmids and oligonucleotides

Diagram of constructs and a list of plasmids and related sequences used in this study are found in **Supplementary Note 1**; LbCpf1 target sites and multiplex crRNA sequences, and SpCas9 target sites are found **Supplementary Table 1**. Plasmids dLbCpf1-p65 (JG1202) and dLbCpf1-VPR (JG1211) were constructed by cloning p65 and VPR domains into dLbCpf1(D832A) (MMW1578) using BstZ17I and Not I sites through isothermal assembly. VPR was amplified from SP-dCas9-VPR which was a gift from George Church (Addgene plasmid # 63798)^19^. Plasmids encoding dLbCpf1-DmrA(x1), dLbCpf1-DmrA(x2), dLbCpf1-DmrA(x3), and dLbCpf1-DmrA(x4) (JG674, JG676, JG693, and YET1000, respectively) were generated by subcloning dLbCpf1(D83A) into AgeI and XhoI digested constructs that have different numbers of DmrA domains (BPK1019, BPK1033, BPK1140, BPK1179 for dSpCas9-DmrA(x1) to dSpCas9-DmrA(X4), respectively) using isothermal assembly. Plasmids encoding dSpCas9(D10A/H840A) effector fusions to VP64, p65, or DmrA were cloned via isothermal assembly (for a complete list of plasmids, please see Supplementary Note). The DmrC entry vector was digested with NruI, and p65 or VPR were inserted via isothermal assembly to generate DmrC-p65 (BPK1169) and DmrC-VPR (MMW948). Single LbCpf1 crRNA expression plasmids were constructed by ligating annealed oligo duplexes into BsmBI-digested BPK3082 (Addgene #78742)^20^. Multiplex LbCpf1 crRNA plasmids were assembled by annealing, phosphorylating, and ligating three pairs of oligonucleotides into BsmBI and HindIII-digested BPK3082, in one reaction (sequences for all oligo pairs are listed in **Supplementary Table 1**).

### Human cell culture and transfection

HEK293 cells were grown at 37° C, in 5% CO2 in Dulbecco’s Modified Eagle Medium (DMEM) with 10% fetal bovine serum and 1% penicillin and streptomycin. Media supernatant was analyzed biweekly for the presence of Mycoplasma. For the GFP activation assays in Supplementary Fig. 1, 500 ng of plasmid DNA expressing dCas9-VPR, dAsCpf1-VPR or dLbCpf1-VPR and 500 ng of crRNA plasmid were co-transfected into HEK293 cells that stably encode GFP under a Tet-On 3G doxycycline-inducible promoter (Takara Clontech). Median GFP fluorescence was measured 3 days post-transfection using a LSR II Flow Cytometer (BD) and values were normalized to HEK293-GFP cells with no plasmids transfected. For LbCpf1 experiments on endogenous genes, 750 ng of dLbCpf1-p65 or dLbCpf1-VPR plasmids with 250 ng of LbCpf1 crRNA plasmids were co-transfected using a 3ul of TransIT^®^-LT1 Transfection Reagent (Mirus, cat# MIR2300) into HEK293 cells in a 12-well plate. For experiments with dLbCpf1 fusions to DmrA domains, 400 ng of dLbCpf1-DmrA fusions plasmids, 200 ng of DmrCp65 or DmrC-VPR plasmids, and 400 ng of LbCpf1 crRNA plasmids were co-transfected using 3 ul of LT1 into HEK293 cells in a 12-well plate. A complete media containing 500 μM A/C heterodimerizer (Takara Clontech) was used at the time of transfection.

### Quantitative reverse transcription PCR

Total RNA was extracted from the transfected cells 72 hours post-transfection using the NucleoSpin^®^ RNA Plus (Clontech, cat# 740984.250), and 250 ng of purified RNA was used for cDNA synthesis using High-Capacity RNA-cDNA kit (ThermoFisher, cat# 4387406). cDNA was diluted 1:20 and 3 ul of cDNA was used for quantitative PCR (qPCR). qPCR reaction samples were prepared using cDNA, SYBR (ThermoFisher, cat# 4385612), and primers detecting each target transcript. Primer sequences are listed in Supplementary Table 2. qPCR was performed using Roche LightCycler480 with the following cycling protocols (**Supplementary Table 2**). When Ct values are over 35, we considered them as 35, because Ct values fluctuate for very low expressed transcripts. Samples that were transfected with LbCpf1 crRNA backbone plasmid, BPK3082 were used as negative controls, and the levels of fold activation over negative controls were normalized to the expression of *HPRT1*.

## VEGFA ELISA

HEK293 cells were seeded in 24-well plates roughly 20 hours prior to transfection using Lipofectamine 3000 (Thermo Fisher Scientific). Unless otherwise indicated, 250 ng of dCas9-DmrA plasmid was co-transfected with 250 ng of sgRNA plasmid (single VEGFA site 3 sgRNA or pooled VEGFA sites 1, 2, and 3 sgRNAs^1^) and 125 ng of DmrC-effector plasmid. A complete media exchange was performed approximately 20 hours post-transfection, with the exchanged media containing 500 μM A/C heterodimerizer (Takara Clontech), unless otherwise noted. Approximately 42 hours post-transfection, supernatant media was removed and clarified prior to analysis using a Human VEGF Quantikine ELISA kit (R&D Systems). Optical density of stopped ELISA reactions was determined using a Model 860 Microplate reader (Bio-Rad).

## SUPPLEMENTARY FIGURE LEGENDS

**Supplementary Figure 1. Engineering dCpf1 for gene**

a. A doxycycline-inducible GFP reporter stably integrated into HEK293 cells used to assess CRISPRa activity. Each guide has seven potential target sites.
b. Diagram of dCas9, dLbCpf1, and dAsCpf1 fused to the VP64-p65-Rta (VPR) activator system.
c. Quantification of GFP fluorescence activation 3 days post-transfection with vectors expressing the protein fusions and guide RNAs. The rTTA+Dox serves as a positive control for GFP activation. Fluorescence values were normalized to HEK293-GFP reporter cells without plasmids transfected (lane 1). Each bar is an average of two replicates.

**Supplementary Figure 2. Optimization of a drug-dependent dCas9 activator platform**

a. Titrations of A/C heterodimerizer (left panel) and DmrC-VP64 or DmrC-p65 effector (right panel), using either one or three sgRNAs targeted upstream of the VEGFA transcription start site. Gene activation assessed by VEGFA ELISA; error bars represent s.e.m. for n = 3.
b. Relative differences in gene activation between direct fusions to dCas9 compared to drug-dependent recruitment using DmrA and DmrC fusions, using either one or three sgRNAs targeted upstream of the VEGFA transcription start site. Gene activation assessed by VEGFA ELISA; error bars represent s.e.m. for n = 2, otherwise n = 1.
c. Differences in VEGFA production based on DmrA repeat variation on dCas9-DmrA or DmrA-dCas9, from 1 to 4 copies, using three sgRNAs targeted upstream of the VEGFA transcription start site. Gene activation assessed by VEGFA ELISA; error bars represent s.e.m. for n = 3.

## ACKNOWLEDGEMENTS

We thank Moira Welch and Alexander Sousa for technical assistance with constructing certain plasmids. B.P.K. acknowledges support from Banting (Natural Sciences and Engineering Research Council of Canada) and Charles A. King Trust Postdoctoral Fellowships. This work was supported by the National Institutes of Health R35 GM118158 (J.K.J.), NIH R01 GM107427 (J.K.J.), R01 DA036858 (J.S.W.) and U01 CA168370 (J.S.W.), the Howard Hughes Medical Institute (J.S.W.), and the Desmond and Ann Heathwood Massachusetts General Hospital Research Scholar Award (J.K.J.).

## AUTHOR CONTRIBUTIONS

Y.G.T., B.P.K., J.K.N., J.S.W., and J.K.J. conceived of and designed experiments. Y.G.T., B.P.K., J.K.N., J.H., and J.G. performed experiments. Y.G.T., B.P.K., J.K.N., J.S.W., and J.K.J. wrote the manuscript.

## DATA AVAILABILITY STATEMENT

The data that support the findings of this study are available from the corresponding author upon reasonable request.

## REFERENCES

1. Maeder, M.L. et al. CRISPR RNA-guided activation of endogenous human genes. Nat Methods 10, 977–979 (2013).

2. Perez-Pinera, P. et al. RNA-guided gene activation by CRISPR-Cas9-based transcription factors. Nat Methods 10, 973–976 (2013).

3. Gilbert, L.A. et al. Genome-Scale CRISPR-Mediated Control of Gene Repression and Activation. Cell 159, 647–661 (2014).

4. Shen, J.P. et al. Combinatorial CRISPR-Cas9 screens for de novo mapping of genetic interactions. Nature methods (2017).

5. Konermann, S. et al. Genome-scale transcriptional activation by an engineered CRISPR-Cas9 complex. Nature 517, 583–588 (2015).

6. Zetsche, B., Volz, S.E. & Zhang, F. A split-Cas9 architecture for inducible genome editing and transcription modulation. Nature biotechnology 33, 139–142 (2015).

7. Guo, J. et al. An inducible CRISPR-ON system for controllable gene activation in human pluripotent stem cells. Protein & cell (2017).

8. Bao, Z., Jain, S., Jaroenpuntaruk, V. & Zhao, H. Orthogonal Genetic Regulation in Human Cells Using Chemically Induced CRISPR/Cas9 Activators. ACS synthetic biology (2017).

9. Polstein, L.R. & Gersbach, C.A. A light-inducible CRISPR-Cas9 system for control of endogenous gene activation. Nature chemical biology 11, 198–200 (2015).

10. Maji, B. et al. Multidimensional chemical control of CRISPR-Cas9. Nature chemical biology 13, 9–11 (2017).

11. Zetsche, B. et al. Cpf1 Is a Single RNA-Guided Endonuclease of a Class 2 CRISPR-Cas System. Cell 163, 759–771 (2015).

12. Kim, H.K. et al. In vivo high-throughput profiling of CRISPR-Cpf1 activity. Nature methods 14, 153–159 (2017).

13. Gao, L. et al. Engineered Cpf1 variants with altered PAM specificities. Nat Biotechnol (2017).

14. Fonfara, I., Richter, H., Bratovic, M., Le Rhun, A. & Charpentier, E. The CRISPR-associated DNA-cleaving enzyme Cpf1 also processes precursor CRISPR RNA. Nature 532, 517–521 (2016).

15. Zetsche, B. et al. Multiplex gene editing by CRISPR-Cpf1 using a single crRNA array. Nature biotechnology 35, 31–34 (2017).

16. Kim, S.K. et al. Efficient Transcriptional Gene Repression by Type V-A CRISPR-Cpf1 from Eubacterium eligens. ACS synthetic biology (2017).

17. Zhang, X. et al. Multiplex gene regulation by CRISPR-ddCpf1. Cell Discov 3, 17018 (2017).

18. Tang, X. et al. A CRISPR-Cpf1 system for efficient genome editing and transcriptional repression in plants. Nature plants 3, 17018 (2017).

19. Chavez, A. et al. Highly efficient Cas9-mediated transcriptional programming. Nature methods 12, 326–328 (2015).

20. Kleinstiver, B.P. et al. Genome-wide specificities of CRISPR-Cas Cpf1 nucleases in human cells. Nature biotechnology 34, 869–874 (2016).

21. Kim, D. et al. Genome-wide analysis reveals specificities of Cpf1 endonucleases in human cells. Nature biotechnology 34, 863–868 (2016).

22. Chavez, A. et al. Comparison of Cas9 activators in multiple species. Nature methods 13, 563–567 (2016).

23. Rivera, V.M., Berk, L. & Clackson, T. Dimerizer-mediated regulation of gene expression in vivo. Cold Spring Harbor protocols 2012, 821–824 (2012).

24. Banaszynski, L.A., Liu, C.W. & Wandless, T.J. Characterization of the FKBP.rapamycin.FRB ternary complex. Journal of the American Chemical Society 127, 4715–4721 (2005).

25. Han, K. et al. Synergistic drug combinations for cancer identified in a CRISPR screen for pairwise genetic interactions. Nature biotechnology (2017).

26. Adamson, B. et al. A Multiplexed Single-Cell CRISPR Screening Platform Enables Systematic Dissection of the Unfolded Protein Response. Cell 167, 1867–1882.e1821 (2016).

27. Tsai, S.Q. et al. Dimeric CRISPR RNA-guided FokI nucleases for highly specific genome editing. Nat Biotechnol 32, 569–576 (2014).

28. Nissim, L., Perli, S.D., Fridkin, A., Perez-Pinera, P. & Lu, T.K. Multiplexed and programmable regulation of gene networks with an integrated RNA and CRISPR/Cas toolkit in human cells. Molecular cell 54, 698–710 (2014).

29. Xie, K., Minkenberg, B. & Yang, Y. Boosting CRISPR/Cas9 multiplex editing capability with the endogenous tRNA-processing system. Proc Natl Acad Sci U S A 112, 3570–3575 (2015).

30. Xu, L., Zhao, L., Gao, Y., Xu, J. & Han, R. Empower multiplex cell and tissue-specific CRISPR-mediated gene manipulation with self-cleaving ribozymes and tRNA. Nucleic Acids Res (2016).

31. Kabadi, A.M., Ousterout, D.G., Hilton, I.B. & Gersbach, C.A. Multiplex CRISPR/Cas9-based genome engineering from a single lentiviral vector. Nucleic Acids Res 42, e147 (2014).

32. Wong, A.S. et al. Multiplexed barcoded CRISPR-Cas9 screening enabled by CombiGEM. Proc Natl Acad Sci U S A 113, 2544–2549 (2016).

33. Hilton, I.B. et al. Epigenome editing by a CRISPR-Cas9-based acetyltransferase activates genes from promoters and enhancers. Nature biotechnology 33, 510–517 (2015).

34. Liu, X.S. et al. Editing DNA Methylation in the Mammalian Genome. Cell 167, 233–247.e217 (2016).

35. Gao, Y. et al. Complex transcriptional modulation with orthogonal and inducible dCas9 regulators. Nature methods 13, 1043–1049 (2016).

